# The *Medicago truncatula* HKT family: Ion transport properties and regulation of expression upon abiotic stresses and symbiosis

**DOI:** 10.1101/720474

**Authors:** Julien Thouin, Man Yuan Guo, Ikram Zribi, Nicolas Pauly, Mohammed Mouradi, Cherki Ghoulam, Hervé Sentenac, Anne-Aliénor Véry

## Abstract

Soil salinity is one of the most important abiotic stresses affecting plant growth. In legumes, symbiotic nitrogen fixation in nodules is affected by salt stress, and salinity tolerance is variable among species. Genes from the High affinity K^+^ Transporter (HKT) family are known to play crucial roles in salt stress tolerance in different plant species. In legumes these transporters are still very poorly characterized.. Here we study the HKT transporter family from the model legume *Medicago trunacatula*, which is moderately tolerant to salinity. The genome of this species comprises five *HKT* genes, hereafter named *MtHKT1;1* to *MtHKT1;5*. Phylogenetic analysis indicated that the MtHKT polypeptides belong to HKT subfamily 1. Three members (*MtHKT1;2, MtHKT1;4* and *MtHKT1;5*) of the *Medicago truncatula* family were cloned and expressed in *Xenopus* oocytes. Their electrophysiological properties revealed a permeability 10 times higher for Na^+^ than for K^+^ and varying rectification properties. Expression analyses of the three *MtHKT* genes under different biotic and abiotic conditions suggested that MtHKT1;5 is the main transporter from this family in the root, the three genes sharing a decrease of expression in drought and salt stress conditions in non inoculated plants as well as plants inoculated with rhizobia. In the shoot, the three MtHKT would be present at similar levels independently on the applied stresses. Based on biomass and ion content analysis, the nodule appeared as the most sensitive organ to the applied salt and drought stresses. The level of expression of the three *MtHKT* genes was strongly decreased by both stresses in the nodule.

## INTRODUCTION

Soil salinity is one of the most important abiotic stresses affecting plant growth (Munns et Gilliham, 2015). Most crop species are poorly tolerant to salt stress and displays reduced yields when the electrical conductivity of the soil solution (at saturation) becomes higher than ca. 4 dS/m (USDA-ARS, 2008), *i.e.* higher than that of a solution containing *ca.* 40 mM NaCl and, which generates an osmotic pressure of approximately 0.2 MPa (Munns et Tester, 2008; Marschner, 2012). Such soils are then classified as saline. Natural salinity and salinization due to irrigation with poor quality water challenge agriculture in 6 % to 10 % of the earth’s land area (Eynard et al., 2005; Munns et Tester, 2008). Enhanced demand for irrigation due to both population increase and climate change, are predicted to dramatically speed up loss of arable land (Eckholm, 1975; Shabala et al., 2014).

Soil salinity results in both an osmotic and an ionic stress for most plants. Osmotic stress results from the fact that high external salt concentrations reduce soil water availability due to the decrease in water potential, and thus make it harder for roots to extract water, which is a situation resembling drought. The so-called ionic stress corresponds to the fact that high salt concentrations within the plant tissues have deleterious effects on plant metabolism and physiology. Molecular, genetic and physiological analyses indicate that Na^+^ rather than Cl^-^ is the major cause of the toxicity in most cases (Munns et Tester, 2008). Screening of mutant plants and searches for QTLs of plant tolerance to salt stress have identified Na^+^ transporters involved in the control of long distance transport of this cation between roots and shoots via the plant vasculature in the model plant *Arabidopsis thaliana* (Shi et al., 2002; Berthomieu et al., 2003) as well as in e.g. cereals, tomato, grapevine (Ren et al., 2005; Byrt et al., 2007, Asins et al., 2013; Campbell et al., 2017, Henderson et al., 2018).

Salt tolerance involves highly complex mechanisms, from osmolyte and osmoprotectant synthesis to anatomical responses, including specific adaptations in some species, such as succulence or salt excretion by salt glands at the leaf surface. A major determinant of salt tolerance common to all plants is the capacity, under high Na^+^ concentrations, to efficiently control Na^+^ and K^+^ uptake and long distance transport. Amongst the transport systems playing crucial roles in this control are members from the so-called HKT family. The acronym HKT means High affinity K^+^ Transporters (Schachtman et Schroeder, 1994) but these systems are either selective for Na^+^ (transporters belonging to HKT subfamily 1), with respect to all other inorganic monovalent cations, or permeable to both Na^+^ and K^+^ with respect to the other monovalent cations (subfamily 2; see Discussion).

The HKT family is still very poorly characterized in legumes. The variation in salt tolerance in legumes species appears to be very great (Läuchli, 1984) and salinity strongly affects symbiotic nitrogen fixation in legume nodules (Delgado et al., 1994). Here we report on HKT transporters from the model legume *Medicago trunacatula*, which can be found in subarid habitats and exposed to soil salinity and drought conditions (Badri et al., 2007; Gil-Quintana et al., 2015). When compared with other legume crops such as pea, bean and soybean, *M. truncatula* is rather tolerant to soil salinity and drought (Moran et al., 1994; González et al., 1998; Costa França et al., 2000; Gálvez et al., 2005).

## Material and methods

### Plant material and experimental conditions for *MtHKT* expression analysis

*M. truncatula* (ecotype Jemalong A17) seeds were scarified with sulfuric acid (99%) for 10 min, sterilized in 6% sodium hypochlorite solution for 1 min and disposed on inverted 0.8% agar plates. The plates were kept for 48 h in the dark at 4 °C and thereafter transferred at 21 °C during 24 h for germination. Seedlings with a radicle long of about 2 cm were then transferred onto pot filled with vermiculite/sand substrate (1/3, V/V). Plants were grown with a photoperiod of 16 h light (25 °C) and 8 h dark (21 °C) in a growth chamber (70% humidity, 200 μE.m^-2^.s^-1^ light intensity). They were watered with Fahräeus solution (Barker et al., 2006): 0.5 mM MgSO_4_, 0.7 mM KH_2_PO_4_, 0.4 mM NaH_2_PO_4_, 10 μM Fe-EDTA, 1 mM urea and 1 μg.mL^-1^ of each of the following micronutrient: MnSO_4_, CuSO_4_, ZnSO_4_, H_3_BO_3_, and Na_2_MoO_4_. Inoculation, when performed, took place after one week of growth. The rhizobial strain was *S. meliloti* 2011-GFP (Cheng et Walker, 1998). Rhizobia were grown in TY liquid medium (3 g.L^-1^ yeast extract, 5 g.L^-1^ tryptone and 6 mM CaCl_2_) supplemented with 10 μg.l^-1^ of tetracycline until an OD_600_ of 0.8 was reached. The suspension was then centrifuged for 15 min at room temperature. The pellet was resuspended with 100 mL water, this washing step being repeated twice, allowing antibiotics removal. The resuspended cells were diluted to OD_600_ 0.05 in Fahräeus solution, which was used for watering and thereby inoculating plants. The inoculated plants were then watered with Fahräeus solution without urea.

### Determination of Na^+^ contents

Plant material was weighed (for fresh weight) or dried for at least 48 hr at 70 °C and then weighed (for dry weight). Plant material was then incubated in 0.1 N chlorhydric acid for 24 hr for ion extraction. Na^+^ concentrations in the extracts were determined by flame photometry (SpectrAA220 fast sequential atomic absorption spectrometer, Varian, Inc.).

### Cloning of *Medicago truncatula HKTs* cDNA

Total RNA was extracted from 4-week old plants using Qiagen RNeasy Plant Mini kit according to the supplier’s instructions. The RNA was treated with DNase I (Thermo Scientific). Reverse transcription cDNA synthesis was performed with the cDNA First Strand cDNA Synthesis kit (Thermo Scientific) according to the manufacturer’s recommendations. Based on the available *M. truncatula* genome sequence (Krishnakumar et al., 2014; Tang et al., 2014), primers were designed for each of the 5 *HKT* genes identified in this genome (Supplemental Table 1). The amplification protocol, after denaturation of the template cDNA at 94 °C for 1 min, consisted of 35 cycles of 10 sec at 94 °C, 30 sec at 53 °C, and 1 min at 72 °C, with a final extension for 5 min at 72 °C. High Proof Polymerase (Bio-Rad) was used. Amplified products were cloned using the pGEM-T Easy vector system (Promega). Successful isolation of *MtHKT*-type cDNA was confirmed by sequencing and comparison with the genome sequence.

### Analysis of gene expression by real-time PCR

Total RNA was extracted using the Qiagen RNeasy Plant Mini kit according to the supplier’s instructions. Its quality was checked with a NanoDrop ND-1000 spectrophotometer (NanoDrop Technologies, Wilmington, USA). Prior to cDNA synthesis, total RNA was treated with DNase I (Thermo Scientific). Reverse transcription reaction for cDNA synthesis was performed from 1 to 2 μg of total RNA treated with DNAse I and with the cDNA First Strand cDNA Synthesis kit (Thermo Scientific) according to the manufacturer’s recommendations. The cDNA was stored at −20 °C until use.

Real-time PCR (qRT-PCR) analysis of *M. truncatula HKTs* genes was performed using a LightCycler® 480 (Roche Molecular Systems, USA). cDNA amplification was carried out with 40 ng of total cRNA, SYBR® Premix Ex Taq™ II (TAKARA Bio Inc) and specific primers designed with the Primer3web program (http://primer3.ut.ee) (Supplemental Table 1). The reaction was done according to the manufacturer’s recommendations. Three technical replicates were run for each of the three biological replicates. As negative controls, water was used in the reaction mixture instead of cDNA. The temperature profile was 95 °C for 30 sec, followed by 40 cycles of 95 °C for 10 sec, 60 °C for 10 sec, and 72 °C for 15 sec. After each amplification step, fluorescence was measured and at the end of the amplification, a melting curve was generated by continuous reading of fluorescence from 64 °C to 95 °C.

Different housekeeping candidate genes were tested *MtTC77416* (*Medtr1g079510*), *MtTubulin* (*Medtr8g107250*), *MtActin* (*Medtr7g026230*), *MtEF1B* (*Medtr3g058940*). The expression of each *MtHKT* gene was normalized to the expression of the *MtTC77416* gene, which was shown to be of constant expression in different tissues in *Medicago truncatula* (Lohar et al., 2006; Serwatowska et al., 2014) and in our data set. The data obtained were analyzed manually and calculations were done to estimate the absolute expression of each gene.

### Expression in *Xenopus laevis* oocytes and two-electrode voltage-clamp analysis

*MtHKT* cDNA were subcloned into the pGEMXho vector (derived from pGEMDG; D. Becker, Würzburg) downstream from the T7 promoter and between the 5’ and 3’-untranslated regions of *Xenopus β-globin* gene. Capped and polyadenylated copy RNA (cRNA) were synthesised *in vitro*, from linearized vector, using a mMESSAGE mMACHINE T7 kit (Ambion). Oocytes, isolated as described previously (Véry et al., 1995), were injected with either 50 nl of *MtHKT1;2* (15 ng), *MtHKT1;4* (15 ng), *MtHKT1;5* (30 ng) cRNA or 50 nl of water for control oocytes, and then kept at 18°C in ND96 solution (96 mM NaCl, 2 mM KCl, 1.8 mM CaCl_2_, 1 mM MgCl_2_, 2.5 mM sodium pyruvate, and 5 mM HEPES/NaOH, pH 7.4) supplemented with 0.5 mg.l^−1^ of gentamicin, until electrophysiological recordings.

Whole-oocyte currents were recorded using the two-electrode voltage-clamp technique 2 days after cRNA injection, as described by Mian et al. (2011). All electrodes were filled with 3 M KCl. The external solution bathing the oocyte was continuously percolated during the voltage-clamp experiment. All bath solutions contained a background of 6 mM MgCl_2_, 1.8 mM CaCl_2_, and 10 mM MES-1,3-bis[tris(hydroxymethyl)methylamino]propane, pH 5,6. Monovalent cations were added to the background solution as glutamate or chloride salts. D-Mannitol was added to adjust the osmolarity, which was set to 220–240 osmM in every solution. To extract MtHKT-mediated currents from total oocyte currents, mean currents recorded in water-injected control oocytes from the same batch in the same ionic conditions were subtracted from the currents recorded in the HKT-expressing oocytes.

### Multiple alignment and phylogenetic analysis

The polypeptide sequences were first aligned with ClustalO 1.2.1 (Mareuil et al., 2017) and phylogenetic tree was generated with PhyML 3.1 software (http://www.atgc-montpellier.fr/phyml/binaries.php) (substitution model, blossom62) using maximum-likelihood method and 1000 bootstrap replicates.

## Results

### Identification and phylogenetic analysis of *M. truncatula* HKT transporters

Analysis of the *Medicago truncatula* genome database (Krishnakumar et al., 2014) revealed the existence of 5 High-affinity Potassium Transporter sequences, all present in a 50 kb cluster in chromosome 6. These sequences were named *MtHKT1;1* to *MtHKT1;5* according to their relative position along this cluster (Figure 1B). RT-PCR experiments carried out to clone the corresponding cDNAs were successful for 3 genes, *MtHKT1;2, MtHKT1;4* and *MtHKT1;5*. The amplified cDNAs strictly corresponded to the available plant genome sequence, without any difference. No DNA fragment could be amplified for *MtHKT1;1* and *MtHKT1;3*, whatever the plant growth conditions (control, drought stress or salt stress), neither from roots nor from shoots. *In silico* analysis of the genome sequence indicates that *MtHKT1;1* displays an early stop codon in its open reading frame, indicating that this gene cannot encode a typical HKT transporter

Phylogenetic analysis of the HKT families from *M. truncatula*, rice and *Arabidopsis thaliana* (8 members in rice, cv. Nipponbare, and 1 member in Arabidopsis) indicated that the MtHKT polypeptides would belong to subfamily 1, which gathers transporters permeable to Na^+^ and displaying strong selectivity for this cation against K^+^ (Véry et al., 2014). Electrophysiological characterization of the cloned MtHKT transporters was carried out using the *Xenopus* oocyte system.

**Figure 1:**
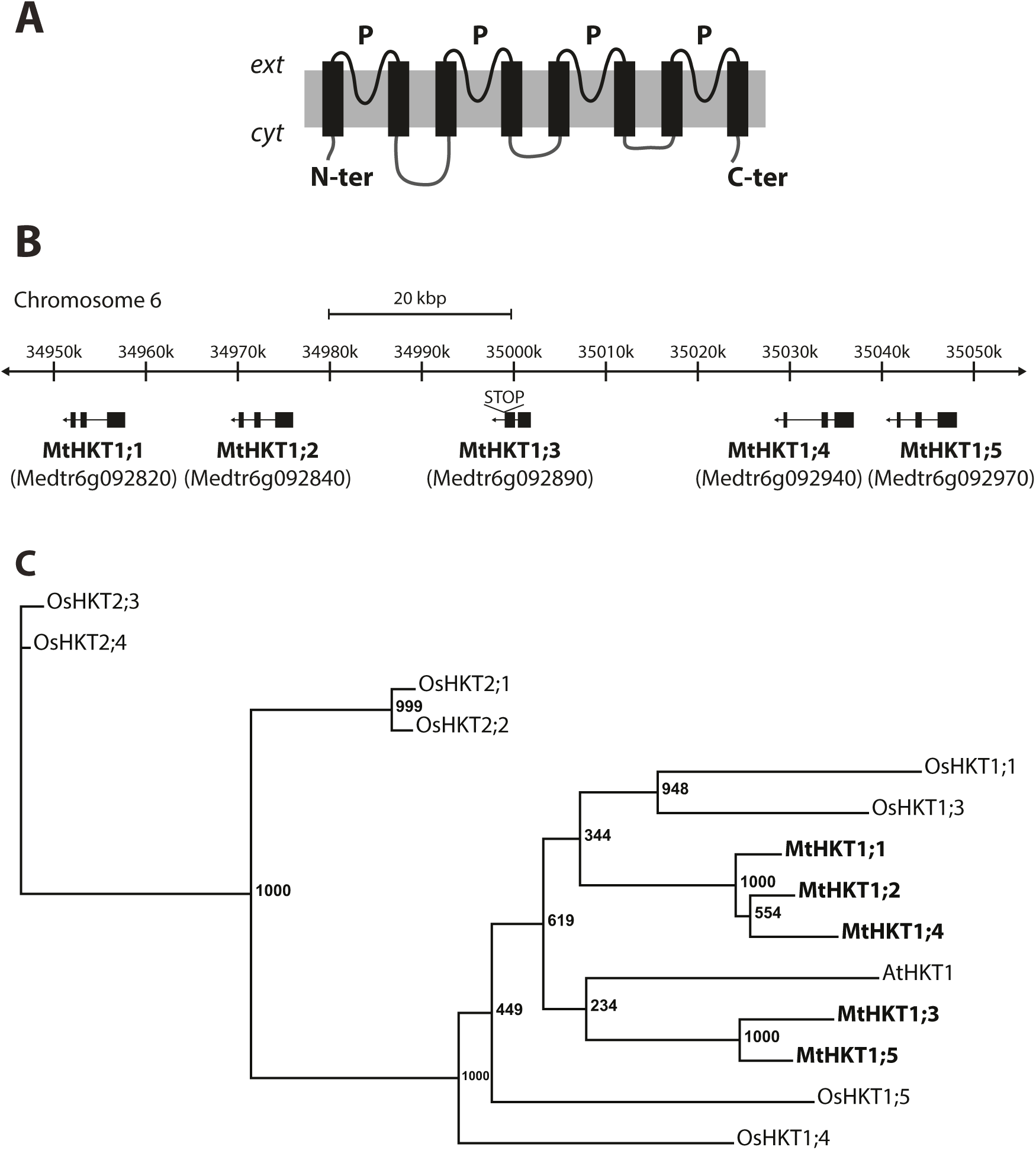
The HKT transporter family of *Medicago truncatula*. **A**: Topology of an HKT transporter based on that defined for bacterial counterparts. N-ter/C-ter, N-terminus/C-terminus of the polypeptide; *ext*/*cyt*, extracellular/cytoplasmic side; P, pore-forming domain; from Véry *et al.*, 2014. **B**: Illustration of annotated portion of chromosome 6 (34,950,000..35,050,000) of the *M. truncatula* genome. The lenght of the representated portion of chromosome is 100 kbp and the position on the chromosome 6 is from 34,950,000 to 35,050,000 base pair. STOP: represent a stop codon on the sequence of MtHKT 1.3. **C**: Phylogenetic relationship of *A. thaliana, O. sativa* and *M. truncatula* HKT proteins. The HKTs polypeptides were aligned with Clustal Omega (https://galaxy.pasteur.fr). The unrooted phylogenic tree was generated with PhyML software (http://www.atgc-montpellier.fr/phyml/binaries.php) using the maximum-likelihood method and 1000 bootstrap replicates. The tree was drawn with FigTree (http://tree.bio.ed.ac.uk/software/figtree/).

### MtHKT conductances upon expression in oocytes

Oocytes were injected with *MtHKT1;2, MtHKT1;4, or MtHKT1;5* cRNA, or injected with water for control oocytes. Figures 2A, 2B and 2C, for MtHKT1;2, MtHKT1;4 and MtHKT1;5, respectively, provide representative current traces recorded in presence of 3 mM Na^+^ and 0.5 mM K^+^ in control oocytes and in *MtHKT* expressing-oocytes. Control oocytes displayed very low current levels in the whole range of imposed membrane potentials, from -165 to -15 mV, when compared with oocytes injected with *MtHKT* cRNA. This indicated that each of the 3 cloned MtHKT transporters was expressed and functional at the oocyte membrane. Current-voltage (I-V) curves obtained from such experiments are displayed by Fig. 2D, 2E and 2F for MtHKT1;2, MtHKT1;4 and MtHKT1;5, respectively, and the corresponding control oocytes. Qualitative differences clearly appear between the 3 MtHKT curves. In the case of MtHKT1;5, the I-V curve is quasi linear in the whole range of recorded currents, inward or outward, as reported for the Arabidopsis AtHKT1 (Xue et al., 2011; Ali et al., 2016) and most other HKT subfamily 1 transporters characterized from dicots (Fairbairn et al., 2000; Almeida et al., 2014a) as well as from monocots (Suzuki et al., 2016). In contrast, in the case of MtHKT1;4, the shape of the I-V curve clearly shows that the ability of this transporter to mediate inward currents is stronger than its ability to mediate outward currents. In other words, this transporter behaves as an inwardly rectifying conductance. The shape of the MtHKT1;2 I-V curve indicates that this transporter too is endowed with inward rectification, but to a lesser extent than MtHKT1;4. Rectification has been rarely reported in HKT transporters so far (Jabnoune et al., 2009; Ali et al., 2016).

**Figure 2:**
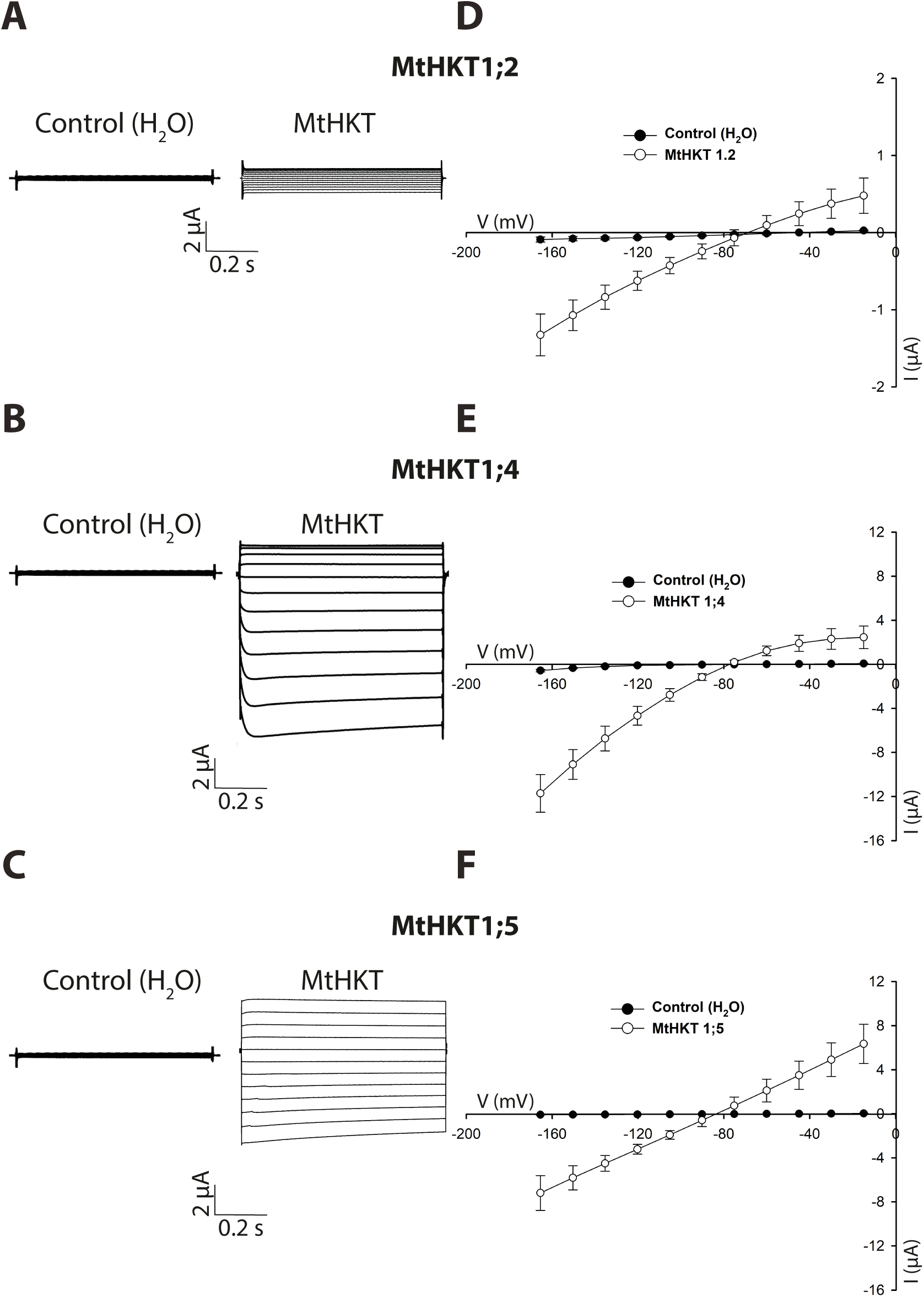
MtHKT expression in *X. laevis* oocytes gives rise to Na^+^ currents. **A, B and C:** current traces recorded in oocytes injected with water or *MtHKT* cRNA, respetiveley *MtHKT1;2* (A), *MtHKT1;4* (B) and *MtHKT1;5* (C), and bathed in 3 mM Na^+^ and 0.3 mM K^+^. **C, D and F:** current-voltage curves in presence of 3 mM Na^+^ and 0.3 mM K^+.^. OoCytes were injected with water or *MtHKT* cRNA, respetiveley *MtHKT1;2* (C), *MtHKT1;4* (D) and *MtHKT1;5* (F). C: Mean±SD, n = 4 and 5 respectively for control and MtHKT1;2. D: Mean±SD, n = 3. C: Mean±SD, n = 5 and 3 respectively for control and MtHKT1;5.

The magnitude of the currents mediated by the three MtHKT cannot be compared directly because the amounts of proteins expressed and functionally targeted to the cell membrane by the oocyte machinery could be different between the three transporters.

### MtHKT are preferentially permeable to Na^+^ when compared to K^+^ and Li^+^

Oocytes were successively bathed in solutions containing either a fixed concentration of K^+^, 0.3 mM, and different concentrations of Na^+^, 0.3, 3 and 10 mM, or a fixed concentration of Na^+^, 0.3 mM, and different concentrations of K^+^, 0.3 or 10 mM. I-V curves obtained for MtHKT1;2, 1;4 and 1;5 in these conditions (and the corresponding I-V curves of the control oocytes) are displayed by Figures 3A, 3B and 3C, respectively. These figures allow to compare, for each MtHKT transporter, the relative positions of the I-V curves obtained in the different ionic conditions and the potentials at which these curves cross the x axis, i.e. the zero current potential, also named reversal potential (Erev). For each MtHKT transporter, Erev was very sensitive to the changes in Na^+^ concentration, being strongly shifted towards more positive values when this concentration was increased. Such sensitivity to the concentration of a given ion demonstrates that the transporter is significantly permeable to this ion. Thus, the 3 MtHKT transporters are permeable to Na^+^. In contrast, the strong increase in K^+^ external concentration, from 0.3 to 10 mM, poorly affected Erev in the 3 MtHKT transporters. The conclusion is thus that these transporters were not significantly permeable to K^+^ in these ionic conditions, where both Na^+^ and K^+^ were simultaneously present in the external solution.

**Figure 3:**
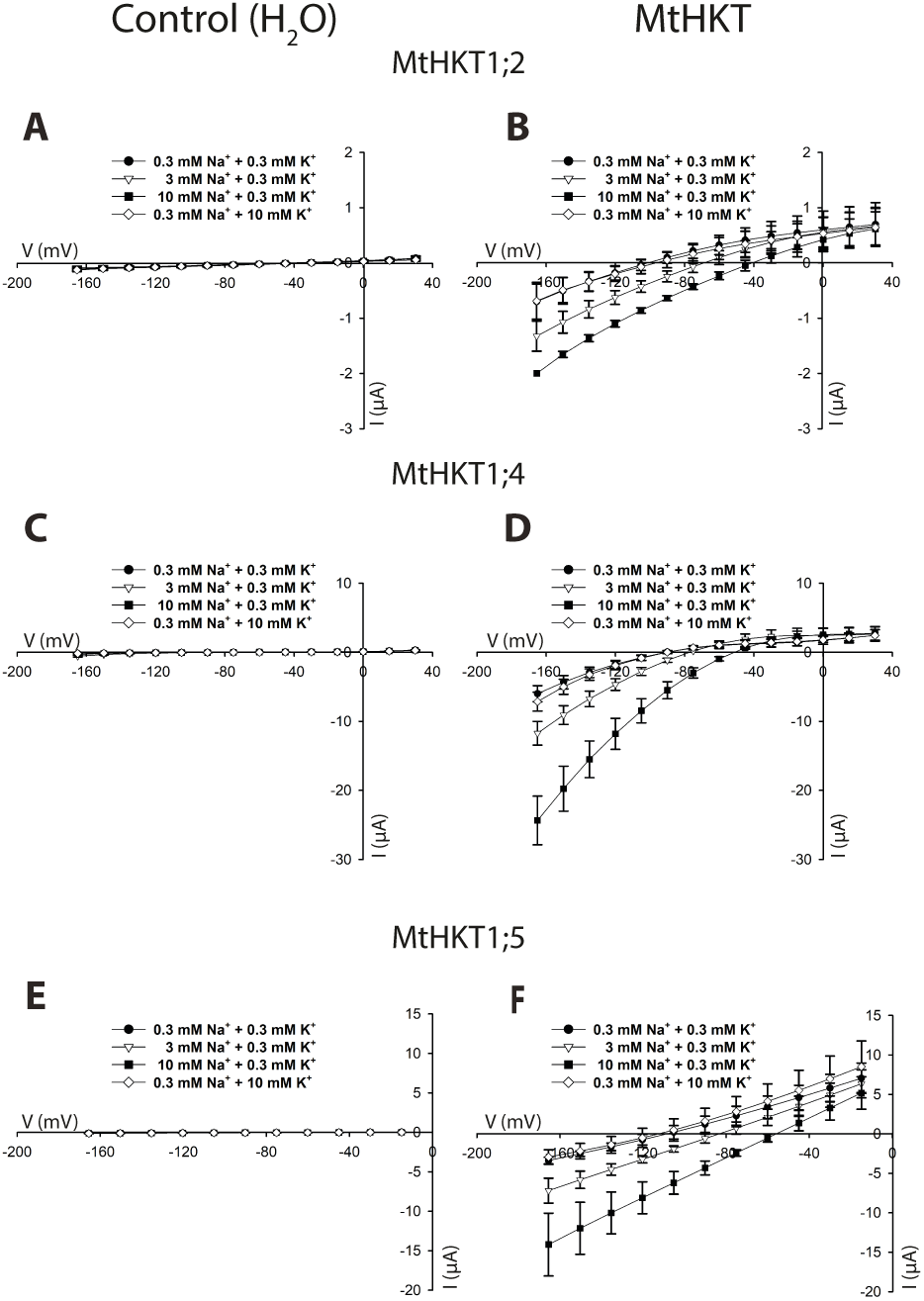
Current-voltage curves in prescence of various K^+^ and Na^+^ concentrations. Oocytes were injected with 50 nl of either water (A, C and E) or *MtHKT1;2* (B), *MtHKT1;4* (D) and *MtHKT1;5* (F) cRNA solution. They were bathed in 4 different solutions, containing variable Na^+^ and K^+^ concentrations, for currents recordings. **A, C and E:** oocytes were injected with water. Mean±SD, n = 5 in A, n = 3 in C and n = 5 in E. **B, D and F:** oocytes were injected with *MtHKT* cRNA. *MtHKT1;2* was injected with 15 ng, *MtHKT1;4* with 15 ng and *MtHKT1;5* with 30 ng. Mean±SD, n = 4 in B, n = 3 in D and F.

MtHKT permeability was then investigated using solutions containing a single monovalent cation, either K^+^, Na^+^ or Li^+^, at a concentration of 10 mM. The corresponding I-V curves are displayed in Figures 4A, 4B and 4C for MtHKT1;2, MtHKT1;4 and MtHKT1;5, respectively. For each of these 3 transporters, Erev and the current magnitude were very similar in presence of K^+^ and Li^+^. The current magnitude was larger and Erev was more positive in presence of Na^+^. These results indicated that K^+^ and Li^+^ were much less permeant than Na^+^ in these ionic conditions. The shift in Erev when K^+^ was replaced by Na^+^ in the external solution was close to +59 mV, +51 mV and +58 mV for MtHKT1;2, MtHKT1;4 and MtHKT1;5, respectively (Figure 4). The “Goldman” equation allows to determine the ratio of the permeability of a transport system to a given ion X, P_X_, to its permeability to another ion Y, P_Y_, from the magnitude of the shift in Erev when X is replaced by Y in the external solution (Dascal et al., 1984). The shifts in Erev determined from Figure 4 when K^+^ is replaced by Na^+^ led to the following values of the P_Na_ to P_k_ permeability ratio: P_Na_/P_K_ = 10.3 for MtHKT1;2, 7.5 for MtHKT1;4 and 9.9 for MtHKT1;5. Thus, the 3 MtHKT transporters were much more permeable to Na^+^ than to K^+^ these ionic conditions too.

**Figure 4:**
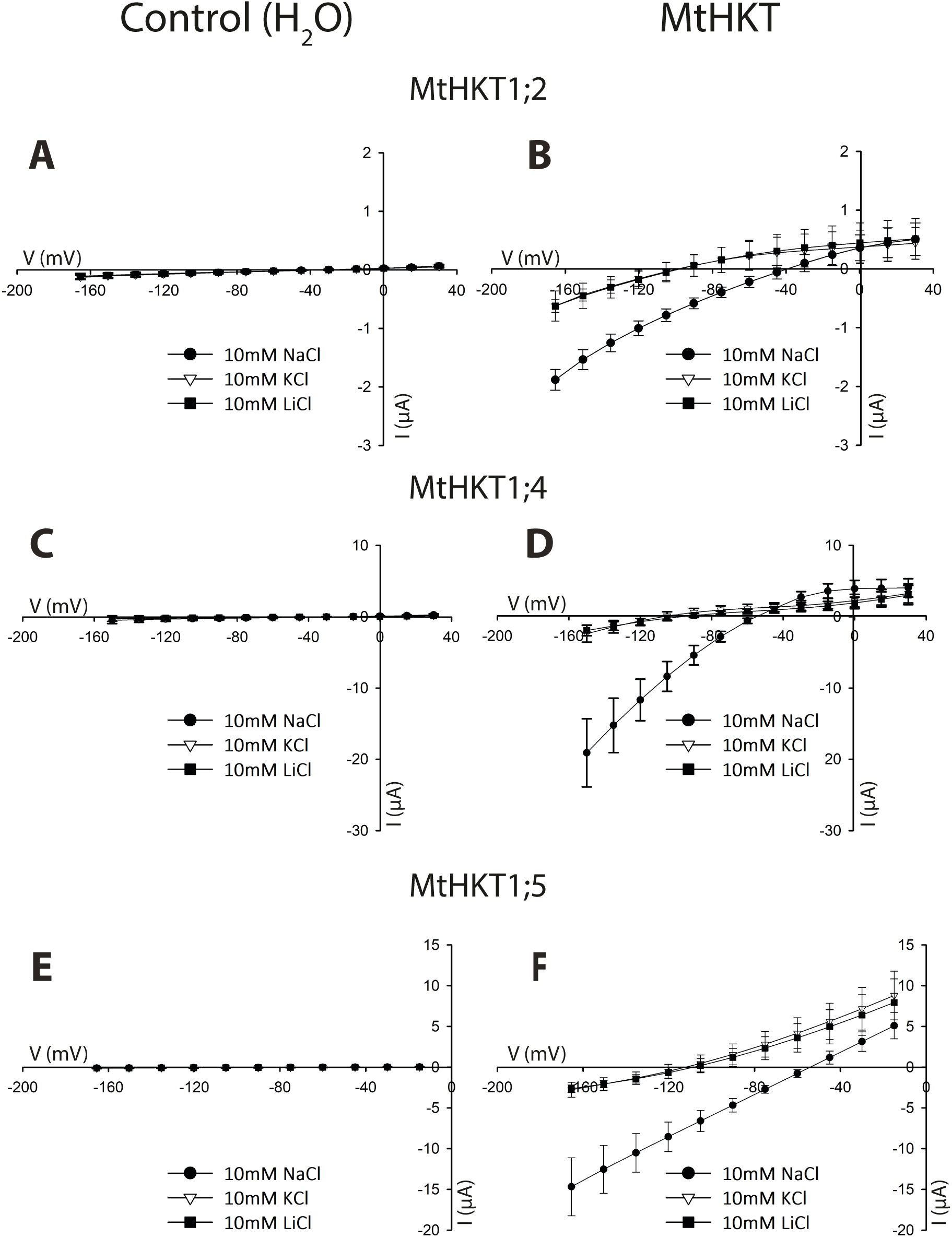
Current-voltage curves in prescence of 10 mM Na^+^, K^+^ or Li^+^. Oocytes were injected with 50 nl of either water (A, C and E) or *MtHKT1;2* (B), *MtHKT1;4* (D) and *MtHKT1;5* (F) cRNA solution. The tested monovalent cations were introduced as chloride salts. **A, C and E:** oocytes were injected with water. Mean±SD, n = 5 in A, n = 3 in C and n = 5 in E. **B, D and F:** oocytes were injected with *MtHKT* cRNA. *MtHKT1;2* was injected with 15 ng, *MtHKT1;4* with 15 ng and *MtHKT1;5* with 30 ng. Mean±SD, n = 3 in B, n = 3 in D and F.

Figure 4B suggests that MtHKT1;4 is endowed with a distinctive functional property, not displayed by MtHKT1;2 and MtHKT1;5 in the same experimental conditions (Figures 4A and 4C). Indeed, the former Figure shows that, at membrane potential more positive than Erev, in other words when an outward current was observed, the magnitude of the current was more important in presence of 10 mM Na^+^ (*i.e.*, “against” 10 mM Na^+^) than in presence of 10 mM K^+^ or Li^+^. Based on the above results, it can be assumed that this current was mostly carried by Na^+^, the most permeant ion in these experimental conditions. Thus, the presence of a high concentration of Na^+^ in the external solution would not reduce the efflux of this ion through MtHKT1;4 but instead would increase this efflux. The simplest hypothesis is that MtHKT1;4 displays a regulatory site present at its extracellular face and with which external Na^+^ ions can interact and stimulate the transporter activity, *i.e.*, its conductance. Within the framework of this hypothesis, the inward conductance of MtHKT1;4 too would display a similar positive regulation by external Na^+^.

### Expression of *MtHKT* genes under different biotic and abiotic conditions

Expression of the 3 *MtHKT* genes in plants inoculated or not inoculated with rhizobia and grown in standard conditions or submitted to drought or salt stress was investigated by Q-RT-PCR (Figure 5). When performed, inoculation was achieved by watering the plants with a solution containing rhizobia (*S. Rhizobium*) 7 days after germination, and the nitrogen source (urea) was then withdrawn from the standard (Fahräeus) nutrient solution. All the plants were 6-week-old when harvested. Plants submitted to drought stress were not watered during the last week of growth, a treatment that was observed to result in wilting symptoms at the end of this last growth week. Plants submitted to salt stress were watered with standard Fahräeus solution supplemented with 100 mM NaCl and thereafter 200 mM NaCl during the last growth week. The wilting symptoms observed at the end of the drought treatment (not shown) and the large increase in plant Na^+^ contents did not significantly affect the final biomass of the plant shoots, roots and nodules (Figure 5A and 5B).

**Figure 5:**
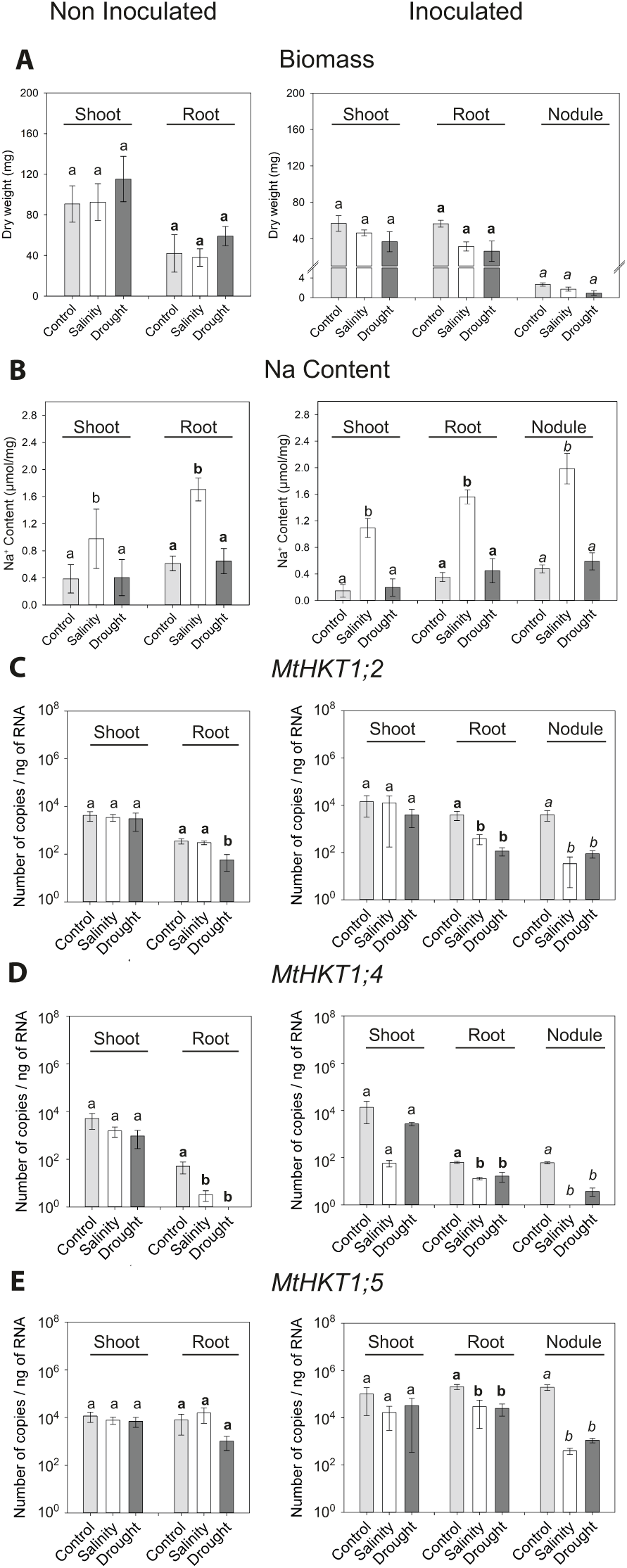
Analysis of *MtHKT* gene expression under salinity and drought stress in *Medicago truncatula* inoculated or non inoculated with rhizobia. Plants were grown for 6 weeks in total in vermiculite/sand soil. They were inoculated or not inoculated with rhizobia at the end of the first week, and submitted or not submitted (control plants) to either salinity or drought stress during the last growth week. For salinity stress, they were watered with 100 mM NaCl the first day of the 6^th^ week, and with 200 mM NaCl two days later. For drought stress, they were not watered during the last 7 days. Roots, shoots and nodules (in inoculated plants) were then harvested for biomass measurements, Na^+^ content assays and Q-RT-PCR analysis of *MtHKT* gene expression. Left panels: non inoculated plants. Right panels: inoculated plants. Light grey, white and dark grey bars: control, salinity and drought treatments. Means ± SD of at least three biological replicates. Statistical comparison was performed within each kind of organs, roots, shoots or nodules, using Turkey’s test. Different letters indicate statistically significant differences between the results from the control, salinity and drought treatments at the level of 5% (Tukey’s test). Normal, **bold** and *italic* letters: comparison of the response in shoot, root and nodules, respectively. **A** : Analysis of dry weight (mg) of non inoculated and inoculated plants. **B**: Analysis of sodium content of non inoculated and inoculated plants. The quantity of sodium (mmol) is normalised by the dry weight (mg). **C, D** and **E**: *MtTC77416* was used as the reference gene for the analysis of the number of copies by Q-RT-PCR experiment. Results are represented with a logarithmic scale. Results of Q-RT-PCR for the expression of MtHKT1;2 (**C**), MtHKT1;4 (**D**), MtHKT1;5 (**E**).

In shoots of non-inoculated plants, the 3 *MtHKT* genes were found to have similar levels of expression (transcript accumulation) and their expression was not significantly sensitive to the drought stress and salt stress (Figure 5C, 5D and 5E). Inoculation tended to increase the level of expression of the three genes in shoots in control conditions but not upon stresses and did not modify significantly the plant sensitivity to the drought and salt stresses (Figure 5A, 5B).

In the roots, the 3 genes differed in their expression levels: in absence of drought and salt stress, the *MtHKT1;5* transcript appeared as the most abundant and the *MtHKT1;4* transcript as the less abundant, whether the roots were inoculated or not. The drought stress in the case of *MtHKT1;2*, and both the drought and salt stresses in the case of *MtHKT1;4*, significantly reduced the level of expression of these genes in roots of non inoculated plants. In inoculated plants, root expression of the 3 genes was reduced by each of the two stresses (Figure 5C, 5D and 5E).

In nodules, the expression levels of the 3 *MtHKT* genes and their variations in response to the drought and salt stresses were similar to those observed in inoculated roots, *i.e.*, in the absence as well as in the presence of drought and salt stress, *MtHKT1;5* displayed the highest levels of transcripts, and *MtHKT1;2* the lowest levels, and the levels of transcripts of each of the 3 genes were significantly reduced by each of the two stresses, drought or salinity (Figure 5C, 5D and 5E).

## Discussion

Phylogenetic analyses have identified 2 subfamilies within plant HKT transporters (Platten et al., 2006), and functional analyses have provided evidence that these 2 subfamilies differ in their K^+^ and/or Na^+^ transport ability. The distinctive property of subfamily 1 members is to be selectively permeable to Na^+^, when compared to K^+^, while subfamily 2 members are significantly permeable to both Na^+^ and K^+^ and have the ability to behave as Na^+^-K^+^ symporters at least when heterologously expressed in animal cells (Rubio et al., 1995; Uozumi et al., 2000; Oomen et al., 2012; Sassi et al., 2012). It should however be noted that some HKT members from subfamily 1 could display a physiologically significant permeability to K^+^ when heterologously expressed in yeast (Ali et al., 2012). Subfamily 1 is present in both dicots and monocots, while members of subfamily 2 have been identified only in monocots so far. Another difference with respect to the HKT family between dicots and monocots can be the family size, which appears to be smaller in dicots. Indeed, when the first genome sequence became available, it appeared that this family comprised a single member in Arabidopsis or poplar, or two members in tomato (Asins et al., 2013), while 8 or 9 *HKT* genes, depending on the cultivar, could be identified in rice, subfamilies 1 and 2 having a similar number of members, 4 or 5. Also, 10 or so *HKT* genes are present in each wheat genome. It should however be noted that recent genome sequences do not support such a dichotomic view since 5 *HKT* genes are present in the dicot *Vitis vinifera* (grape vine).

The number of *HKT* genes seem to be rather variable in legumes. As shown in the present report, *M. truncatula* possesses 5 *HKT* (including one pseudo-gene), which are present within a cluster in chromosome 6. *In silico* analysis (not shown) indicates that the *Lotus japonicus* genome harbors 5 *HKT* genes, forming 2 clusters, the first one located in chromosome 5 and comprising 2 genes, (*Lj5g3v0196960* and *Lj5g3v0196990*), and the second one located in chromosome 2 and comprising 3 genes (*Lj2g3v0914290, Lj2g3v0914260* and Lj2g3v0914190) (Sato et al., 2008). In soybean (*Glycine max*), the *HKT* family displays 4 members, at four different locations in the genome (*Glyma.01g002300, Glyma.06g271600, Glyma.07g130100* and *Glyma.12g133400*) (Schmutz et al., 2010). In contrast, common bean (*Phaseolus vulgaris*) has only two *HKT* genes, (*Phvul.004G177200, Phvul.004G177300*), which form a cluster (Schmutz et al., 2014). The Chickpea (*Cicer arietinum*) possesses one or two *HKT* genes depending of the sequence database (Varshney et al., 2012; Varshney et al., 2013). Altogether, these data indicate that the size of this transporter family in dicots is not as reduced as initially thought and can be rather variable even within a same family.

Based on phylogenetic criteria, the 4 HKT transporters identified in *M. truncatula* belong to HKT subfamily 1. Three from them, whose transcripts could be easily detectable in shoots and roots, have been functionally characterized by electrophysiological analyses in *Xenopus* oocytes. They all appear as typical members from subfamily 1, being much more permeable to Na^+^ than to K^+^, which is consistent with the hypothesis that only HKT subfamily 1 is present in dicots and that the members of this subfamily display strong selectivity for Na^+^ against K^+^. In HKT subfamily 1, a serine residue located in the pore loop of the first MPM domain has been identified as an important determinant of selectivity for Na^+^ (Mäser et al., 2002). MtHKT1;2, MtHKT1;4 and MtHKT1;5 display the serine residue at the corresponding position.

MtHKT1;4 and MtHKT1;1 to a lesser extent display a functional property, rectification in favor of inward currents, that has been rarely reported in HKT transporters (Jabnoune et al., 2009; Ali et al., 2016). The strong inward rectification displayed by TsHKT1;2 from *Thellungiella salsuginea* (Ali et al., 2016), a halophytic *Arabidopsis thaliana* relative (that was previously *Thellungiella halophila*), has been shown to largely result from a single amino acid change, a D in position 207, in the pore loop of the second MPM domain, where AtHKT1 from Arabidopsis displays an N in the corresponding position (position 211 in AtHKT1). Interestingly, while the shape of AtHKT1 current-voltage curves in *Xenopus* oocytes is known to be quasilinear, indicating that the outward and inward transport activity are quite similar, mutation of the N in position 211 to a D results in a marked decrease of the outward currents, clearly highlighting that the mutated transporters is endowed with stronger inward rectification (Ali et al., 2016). It was thus proposed that a D at this position could be a major inducer of rectification in HKT transporter. The three MtHKT we have characterized have an S in the position corresponding to AtHKT1 N211. S is in the same group of amino acids as N, characterized by a polar uncharged side chain. The three MtHKT are however clearly different in terms of rectification capacity. Thus, rectification does not systematically require a D in the position corresponding to AtHKT1 N211 and, in MtHKT transporters, this position does not appear as a major determinant of rectification.

MtHKT1;4 appears to be endowed with another original property, besides rectification. This transporter seems to be activated by external Na^+^, as suggested by Figure 4B. Such activation, together with the inward rectification, could render this transporter especially dedicated to Na^+^ uptake from high Na^+^ concentrations (*e.g.*, in Na^+^ retrieval from the xylem sap?). Conversely, the absence of strong rectification in MtHKT1;2 could allow this transporter to be more specifically involved in Na^+^ secretion (*e.g.*, in Na^+^ release from the phloem vasculature into the root stele and thereby in Na^+^ recirculation from shoots to roots?).

The roles of HKT transporters is probably more documented in monocots than in dicots due to the strong interest that these transporters have for the cereal breeders interested in crop adaptation to soil salinity. Indeed, evidence has been clearly provided that members from HKT subfamily 1 play major roles in the control of Na^+^ shoot contents, and thereby in salt tolerance, in agronomically important cereals such as rice and wheat. For instance, a major salt tolerance QTL in rice, named *SKC1* (shoot K^+^ content) which contains the subfamily 1 transporter *OsHKT1;5* (also named *OsHKT8*) (Lin et al., 2004; Ren et al., 2005), is presently used by plant breeders for increasing salt tolerance in rice cultivars (Ashraf et al., 2012). Major QTL of salt tolerance have been shown to correspond to HKT genes in wheat and maize too (Horie et al., 2009; Zhang et al., 2017). In dicots, the most detailed information is available for the Arabidopsis AtHKT1, which has been shown to be expressed in plant vasculature and to play a major role in salt tolerance by controlling Na^+^ translocation from roots to shoots via the xylem sap and/or Na^+^ recirculation from shoots to roots via the phloem sap (Berthomieu et al., 2003; Horie et al., 2009). Contribution to control of Na^+^ long distance transport in the plant vasculature has also been reported for SlHKT1;2 in tomato (Almeida et al., 2014b), whose gene is located, together with *SlHKT1;1*, in the major tomato QTL involved in Na^+^/K^+^ homeostasis (Asins et al., 2013). In legumes, information on the role of an HKT transporter has been obtained for GmHKT1 from soybean: overexpression of its encoding gene enhances salt tolerance in transgenic tobacco plants and reduces shoot Na^+^ contents (Chen et al., 2011). The transcript levels of the *GmHKT1* gene have been found to be upregulated by salt stress in both roots and leaves of soybean plants (Chen et al., 2011). Such induction of gene expression upon salt stress appears to be the most frequent response in *HKT* genes characterized as playing a role in plant tolerance to salinity. For instance, it has been reported for *AtHKT1* (Wang et al., 2015a), *OsHKT1;1* (Wang et al., 2015b), *OsHKT1;5* (Ren et al., 2005) or *TmHKT1;4-A2* (Tounsi et al., 2016). Induction of *HKT* gene expression upon salt stress is however not a general behavior amongst plants since *SlHKT1;2* from tomato displayed (slightly) reduced expression levels upon a strong increase in external concentration of Na^+^ (from 5 to 70 mM) (Almeida et al., 2014a). A decrease upon salt stress of the expression of *HKT* genes expected to play a role in salt tolerance has also been reported in rice (*OsHKT2;2, OsHKT2;2/1*; Horie et al., 2001; Kader et al., 2006; Oomen et al., 2012) in salt tolerant cultivars. In *M. truncatula*, our results indicate that *MtHKT* gene expression was poorly sensitive to high external Na^+^ concentrations in shoots. The expression was markedly reduced in roots, with the exception of *MtHKT1;2*, and *MtHKT1;5* in non inoculated plants, and in nodules for the 3 *MtHKT* genes. The decrease in expression appeared very strong in nodules, by *ca.* two orders of magnitude, for the 3 *MtHKT* genes. Investigation of the effects of drought stress on the expression of the 3 *MtHKT* revealed changes in transcript accumulation levels that were very similar to those observed upon salt stress. Thus, the osmotic component of salt stress, and not its ionic component, may have underlain the *MtHKT* gene responses to the salt treatment we used. A strong decrease in expression was again observed in nodules for the 3 genes. Symbiotic nitrogen fixation and nodule physiology are strongly sensitive to drought stress (Gil-Quintana et al., 2015; Kunert et al., 2016; Adams et al., 2018). Whether the *MtHKT* gene responses to drought and salt stress we report are merely a consequence of nodule senescence or contributed to nodule adaptation to osmotic/drought stress would be worth to be investigated. Analysis of the expression patterns of the 3 *MtHKT* genes in nodules and their responses to drought and salt stress is under investigation.

## Acknowledgments

This work was supported in part by an ANR grant (ANR-11-BSV7-010-02) (to HS), doctoral grants of the French Ministry of Higher Education and Research (to JT) and of the China Scholarship Council (to M-YG), and a Hubert Curien PRAD (12-09) France-Morocco Partnership.

