## Supplemental Figure 1 for "The *Medicago truncatula* HKT family: Ion transport properties and regulation of expression upon abiotic stresses and symbiosis"

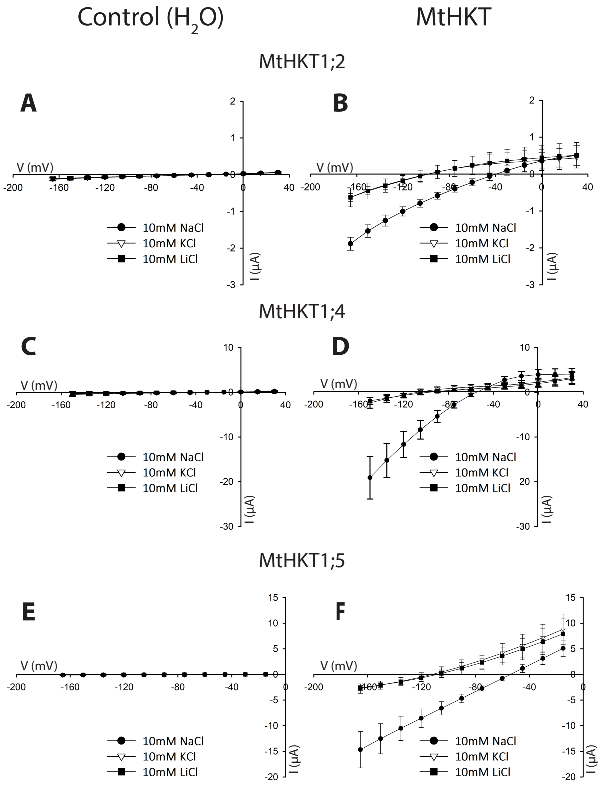


**Figure 4:** Current-voltage curves in prescence of 10 mM $Na^+$, $K^+$ or $Li^+$. Oocytes were injected with 50 nl of either water (A, C and E) or *MtHKT1;2* (B), *MtHKT1;4* (D) and *MtHKT1;5* (F) cRNA solution. The tested monovalent cations were introduced as chloride salts.

**A, C and E:** oocytes were injected with water. Mean±SD, n = 5 in A, n = 3 in C and n = 5 in E.

**B, D and F:** oocytes were injected with *MtHKT* cRNA. *MtHKT1;2* was injected with 15 ng, *MtHKT1;4* with 15 ng and *MtHKT1;5* with 30 ng. Mean±SD, n = 3 in B, n = 3 in D and F.

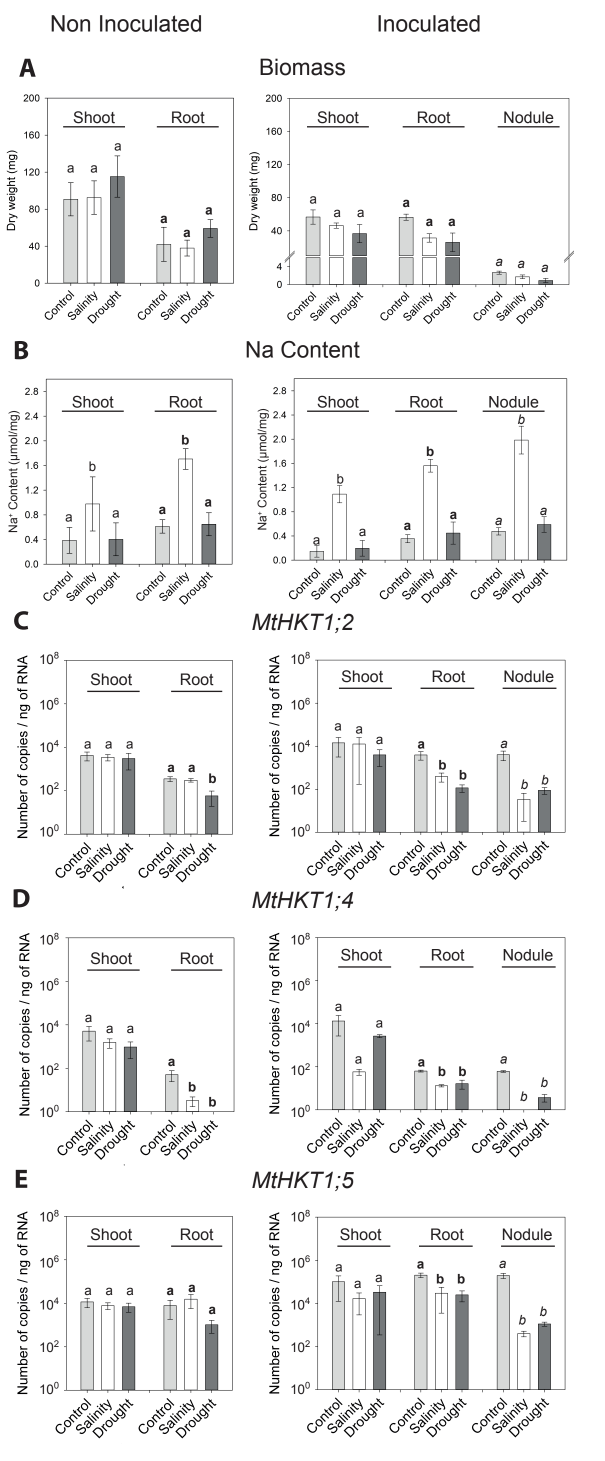


**Figure 5:** Analysis of *MtHKT* gene expression under salinity and drought stress in *Medicago truncatula* inoculated or non inoculated with rhizobia. Plants were grown for 6 weeks in total in vermiculite/sand soil. They were inoculated or not inoculated with rhizobia at the end of the first week, and submitted or not submitted (control plants) to either salinity or drought stress during the last growth week. For salinity stress, they were watered with 100 mM NaCl the first day of the 6th week, and with 200 mM NaCl two days later. For drought stress, they were not watered during the last 7 days. Roots, shoots and nodules (in inoculated plants) were then harvested for biomass measurements, $Na^+$ content assays and Q-RT-PCR analysis of *MtHKT* gene expression. Left panels: non inoculated plants. Right panels: inoculated plants. Light grey, white and dark grey bars: control, salinity and drought treatments. Means ± SD of at least three biological replicates. Statistical comparison was performed within each kind of organs, roots, shoots or nodules, using Turkey's test. Different letters indicate statistically significant differences between the results from the control, salinity and drought treatments at the level of 5% (Tukey's test). Normal, **bold** and *italic* letters: comparison of the response in shoot, root and nodules, respectively.

**A** : Analysis of dry weight (mg) of non inoculated and inoculated plants.

**B**: Analysis of sodium content of non inoculated and inoculated plants. The quantity of sodium (mmol) is normalised by the dry weight (mg).

**C, D** and **E**: *MtTC77416* was used as the reference gene for the analysis of the number of copies by Q-RT-PCR experiment. Results are represented with a logarithmic scale. Results of Q-RT-PCR for the expression of MtHKT1;2 (**C**), MtHKT1;4 (**D**), MtHKT1;5 (**E**).

| | | | $P_A$ ↓ | |
|---|---|---|---|---|
| AtHKT1 | 47 | TTSRPH | DFDL FFTSVSAITV SSM | STVDMEV |
| MtHKT1;2 | 72 | G-ETPK | NLDL FFTSISATTV SSM | STVEMQN |
| MtHKT1;4 | 72 | YQTSPK | NLDL FFTSVSSTTV SSM | STVEMEF |
| MtHKT1;5 | 78 | TSVKPK | DFDL FYTSVSASTV SSM | TSIEMEV |
| TsHKT1;2 | 47 | TTSRPH | DLDL FFTSVSAITV SSM | STIDMEV |

| | | | $P_B$ | |
|---|---|---|---|---|
| AtHKT1 | 188 | DVLSSKEIS | P LTFSVFTTVS TFANCGFV | PT |
| MtHKT1;2 | 239 | QILKNKGLK | M FTFSVFTIVS TFASCGFV | PT |
| MtHKT1;4 | 233 | QVLENKGLK | M FTFSLFTIVS TFSSCGFI | PT |
| MtHKT1;5 | 232 | NILQNKGIN | I ETFSLFTIVS TFASCGYI | PT |
| TsHKT1;2 | 184 | DVLSSKKIS | P LTFSVFTAVS TLSDCGFV | PT |
| | | | * | |

**Supplemental Figure 1: Pore-loop A and B of HKT proteins.**

Sequence comparison of HKT homologs from *A. thalina*, *M. truncatula* and *T. salsuginea*. Amino-acid sequences in the first (PA) and second pore-loop region (PB) are aligned with the use of Clustal Omega (https://galaxy.pasteur.fr). The conserved serine residues in the $P_A$ region are indicated by an arrow (Mäser *et al.,* 2002) and the single amino acid change is indicated by an asterisk, a D in $P_B$ region of *T. salsuginea* (Ali *et al.,* 2016) are highlighted in red.
