## Supplemental Table 1 for "The *Medicago truncatula* HKT family: Ion transport properties and regulation of expression upon abiotic stresses and symbiosis"

**Table1: Q-RT-PCR Primers**

| Gene | Name | Sequence |
|---|---|---|
| *MtTC77416* (*Medtr1g079510*) | TC77416 F | ACGCGCTAATGATTCCAGTG |
| | TC77416 R | AGTGTTGCTGTGTTCCAACG |
| | TC77416GE F | GTTGGCTCATCCGACAGCTC |
| | TC77416GE R | GGACGAAGCTCAATGGCAATC |
| *MtTubulin* (*Medtr8g107250*) | MtTubbQP F | TGCCGAGCTCATTGACTTTG |
| | MtTubbQP R | GTGAGCATCATTCGATCAGGG |
| | MtTubb375 F | TGGTGCCGGAAACAATTTCG |
| | MtTubb375 R | GGTGTGGTGAGTTTGAGGGT |
| *MtActin* (*Medtr7g026230*) | MtActin F | CCGTTCTTTCACTGTACGCC |
| | MtActin R | GGCATGTGGAAGGGCATAAC |
| | MtActin352 F | ACAATGAGCTTCGTGTCGCC |
| | MtActin352 R | ATTTCTCGCTCTGCTGAGGT |
| *MtEF1B* (*Medtr3g058940*) | EF1B F | ATGGTCCCGTCTTTGAGAGC |
| | EF1B R | ATCCATTGTTCAACGTGGGC |
| | EF1B377 F | CTGCATGCTCAGAAGGGAAAC |
| | EF1B377 R | TCTTGACCAAGAGTTGAGCGA |
| *MtHKT1;2* (*Medtr6g092840*) | MtHKT1;2 F1227 | GTACCTTCCTCCTTACACCTCATTCC |
| | MtHKT1;2 R1655 | AATCAGAGAAGTTTCCATGGCTTG |
| | MtHKT1;2 F | TCTAGTGTGCATAACTGAGAGGA |
| | MtHKT1;2 R | CCCTCATCACTCCATTTCCC |
| *MtHKT1;4* (*Medtr6g092940*) | MtHKT1;4 F1209 | GTACCTTCCACCTTACACCTCATTC |
| | MtHKT1;4 R1649 | ATATTCAAGAATCAGAGAAGTTTCCATG |
| | MtHKT1;4 F | TGTCGGAAAGCAGTGAGAGA |
| | MtHKT1;4 R | TCGATGCAACTGTCGTTCAC |
| *MtHKT1;5* (*Medtr6g092970*) | MtHKT1;5 F1272 | GTATCTTCCACCGTACACAACATTTT |
| | MtHKT1;5 R1677 | CTAGGACAGGTGCCATGCCTT |
| | MtHKT1;5 F | TCCCCTCAACTTCAACGTCT |
| | MtHKT1;5 R | AGCTTCCCTTCTGTACTCCA |
